# Reciprocal roles of crowding and serial dependence on visual perception

**DOI:** 10.1101/2025.01.16.633480

**Authors:** John Cass, Robert Jertberg, Erik Van der Burg

## Abstract

Visual perception arises from the interplay between current and prior sensory inputs. Two perceptual phenomena-serial dependence and visual crowding-result from the mandatory integration of retinal information across time and space, respectively. This study investigated, for the first time, their functional relationship for orientation and brightness discrimination tasks. Participants performed orientation and brightness discrimination tasks (blocked) on peripheral targets surrounded by distractors of varying color, orientation, and proximity. Both serial dependence (the influence of prior stimuli on current judgments) and visual crowding (impaired peripheral recognition caused by nearby distractors) were observed. Similar double dissociations in task and stimulus specificity emerged for both phenomena, suggesting functional links mediated by distinct processing mechanisms for brightness and orientation judgments. Additionally, the two effects interacted: crowding reduced serial dependence on subsequent trials, indicating that spatial information is prioritized over temporal information when spatial redundancy is high. Conversely, serial dependence increased crowding for orientation judgments, while for brightness judgments, serial dependence operated additively without affecting crowding. These interactions were spatially specific, occurring only when targets appeared in the same location across consecutive trials. Notably, a positive correlation between crowding and serial dependence was found exclusively for orientation judgments and only when targets appeared at the same location across trials. The reciprocal interaction and positive correlation between serial dependence and crowding observed for orientation judgments indicate that that these processes are intrinsically linked, possibly via a common neural mechanism. In contrast, for brightness judgments, the unilateral effect of crowding on serial dependence—along with the absence of correlation-indicates distinct neural mechanisms. Here, brightness crowding likely precedes serial dependence, reflecting a prioritization of spatial redundancy minimization over temporal stabilization. These findings suggest that the brain strategically integrates spatial and temporal information through distinct yet overlapping mechanisms, balancing spatial precision with temporal stability based on task demands and stimulus characteristics.

## Introduction

Our perception of the visual environment is influenced not solely by immediate retinal input but also by prior visual experiences. Visual aftereffects are a classic manifestation of this historic influence in which prolonged exposure to a visual stimulus induces temporary loss of sensitivity and perceptual distortions in the perceived retinal image (Blakemore & Campbell, 1969; Gibson & Radner, 1937; Maffei et al., 1973; Mollon, 1982; Rhodes et al., 2003; Thompson & Burr, 2009), a phenomenon known as adaptation. More recently, a new short-term form of adaptation known as serial dependence was discovered, differing from classical prolonged adaptation (Fischer & Whitney, 2014; Van der Burg et al., 2013). In Fischer and Whitney’s (2014) study, participants judged the orientation of a series of randomly oriented Gabor patches. Unlike the sensory suppression and perceptual repulsion observed in studies using prolonged adaptation procedures (e.g., the tilt aftereffect), they found that participants’ orientation judgments were systematically biased toward the orientation of the stimulus presented in the previous trial at the same visual location; a phenomenon known as serial dependence. This *attractive serial dependence* (a.k.a. *positive serial dependence*) has since been replicated using various visual stimuli (Liberman et al., 2014; Taubert et al., 2016), tasks (Van der Burg et al., 2019; Van der Burg, Toet, Abbasi, et al., 2021), and sensory modalities (Van der Burg, Baart, et al., 2024; Van der Burg, Toet, Brouwer, et al., 2021).

Attractive serial dependencies are thought to arise from the perceptual assimilation of neural responses evoked by temporally adjacent stimuli. For instance, in the orientation domain, temporal integration across trials leads to a stronger orientation-tuned response when a Gabor stimulus is followed by another with a similar orientation. However, a more complex picture of serial dependencies has emerged, with repulsive effects also being observed. In these cases, perceptual biases shift away from the feature space occupied by the target in the previous trial (Manassi et al., 2023). Although *repulsive serial dependencies* are smaller in magnitude and often less reliable than attractive effects, they are often tightly tuned to specific retinotopic locations (but see (Fritsche et al., 2017)). Various factors, including uncertainty, task, decisional bias, attention, and individual differences, have been found to influence the direction (i.e., assimilation and repulsion) and magnitude of these serial dependencies (Alais et al., 2017; Van der Burg, Baart, et al., 2024) (see Manassi et al. (2023) for a review of visual perception studies). Moreover, some studies report assimilative as well as repulsive effects within a single experiment/study, with the direction governed by whether participants respond to the stimuli (Alais et al., 2017; Van der Burg, Baart, et al., 2024).

Although serial dependencies introduce perceptual distortions (i.e., the percept deviates from reality), including attraction and repulsion effects, they may confer functional advantages. Serial dependencies are thought to reflect a predictive strategy by which the visual system leverages temporal redundancies in the environment. Through this mechanism, attractive serial dependencies enhance the weight of stable visual information in the perceptual array (Fischer & Whitney, 2014), whereas repulsive serial dependencies reflect increased sensitivity to visual changes. This predictive strategy has parallels in spatial visual perception, as illustrated by the phenomenon of visual crowding (Bouma, 1970; Pelli & Tillman, 2008; Whitney & Levi, 2011).

Visual crowding occurs when the identification of a peripheral (target) object is impaired by nearby clutter (flankers) (Bouma, 1970). Like serial dependence, crowding is thought to result from compulsory integration of adjacent (spatially, in this case) target and flanker features, which distorts or disrupts perceptual access to the target’s identity (Dakin et al., 2010; Greenwood et al., 2010; Parkes et al., 2001). Crowding effects are particularly pronounced when target and flankers share similar features (like color & orientation) (Kennedy & Whitaker, 2010; Kooi et al., 1994; Parkes et al., 2001) which introduces spatial redundancy in the retinal image.

While serial dependence and visual crowding have traditionally been studied independently, recent findings suggest they may represent complementary predictive coding mechanisms that enable the brain to exploit statistical redundancies in the visual input, both temporally and spatially (Cicchini et al., 2022). Specifically, the brain appears to average redundant visual information—spatially in the case of crowding and temporally in the case of (attractive) serial dependence—thereby enhancing processing efficiency in line with predictive coding principles (Friston & Kiebel, 2009). Beyond efficiency, retaining a stable perceptual representation of objects while remaining sensitive to change depends upon integrating inputs over space and time specifically when they are likely to come from a common source, a predictive problem for the brain. Stimuli that are temporally and spatially proximal (as well as sharing similar visual features) are likely to fit this description. As such, visual crowding and attractive serial dependence could serve an analogous functional role, both in the reduction of prediction errors and the associated behavioral optimization.

Accordingly, predictive coding offers a compelling explanatory framework for understanding both serial dependencies as well as visual crowding. By down-regulating its response to redundant sensory information, the brain reduces metabolic expenditure and increases efficiency and perceptual reliability. However, predictive coding is not without its limitations, particularly its lack of falsifiability—multiple perceptual outcomes can be explained by adjusting the supposed ’goal’ of the brain. For instance, both repulsive and attractive effects, despite their opposing perceptual outcomes, can be argued to confer distinct adaptive advantages. This adaptability, while theoretically useful, makes predictive coding challenging to test.

In this study, we take a novel approach to investigating the functional relationship between serial dependence and crowding on three levels. First, we explore whether these two phenomena share functional similarities by examining their stimulus and task dependencies. We assess target performance in two tasks: orientation and brightness discrimination judgements, each characterized by distinct feature-specific contingencies. Our previous work (Cass & Van der Burg, 2023) demonstrates that while both tasks are impacted by crowding, the features involved differ, presenting a task-stimulus double dissociation. Specifically, brightness discrimination is modulated by the color similarity between the target and flankers and the target-flanker distance, regardless of their orientation. Conversely, orientation judgments depend on the relative orientation of the target and flankers and the target-flanker distance, independent of color similarity. Here, we aim to determine whether serial dependence exhibits similar stimulus and task specificities to those found in crowding, using identical stimuli for both brightness and orientation tasks. Specifically, we hypothesize that if serial dependence and crowding are functionally related, then we ought to observe a double dissociation between the perceptual task and the physical properties of the target object wherein serial dependencies for brightness judgments on a given trial *t* will be unaffected by the relative orientation of temporally adjacent stimuli (i.e., trial *t*-1), and serial dependencies for orientation judgments will be unaffected by the relative brightness of temporally adjacent stimuli.

Next, we assess whether the strength of crowding—manipulated by varying target-flanker proximity and task-specific target-flanker similarity—on either the current or previous trial influences the magnitude of serial dependence in brightness and/or orientation judgments. Conversely, we investigate whether the observed magnitude of serial dependence modulates the extent and specificity of visual crowding. Not only does this allow for a direct comparison of the functional characteristics of visual crowding and serial dependence phenomena, it also enables us to assess whether the mechanisms underlying visual crowding and serial dependence function independently or interact. If interactions are present, given the distinct stimulus contingencies associated with crowding in brightness and orientation judgments, we can identify the nature of any reciprocal effects between serial dependency and crowding.

Finally, we investigate whether the magnitude of visual crowding and serial dependence effects are correlated within individuals and whether these potential correlations differ according to the task at hand (orientation or brightness judgments). This provides insight into the potential relationship between the phenomena and their task dependencies. A significant correlation would imply that these processes are intimately connected, with their proposed functional similarities perhaps even being rooted in shared underlying mechanisms. With this tripartite approach, we seek to elucidate the nature of the relationship between visual crowding and serial dependence, from the functional similarities they demonstrate, to the way the interact with each other, and finally to the extent to which they co-vary within individuals.

## Method

For the present study, we reanalysed the data from our previously published study (Cass & Van der Burg, 2023).

### Participants

Thirty participants participated in the experiment (vastly exceeding the number typically used in most studies investigating crowding and serial dependencies). Participants were paid €8 per hour and were naive as to the purpose of the experiment. All had normal or corrected-to-normal vision. Informed consent was obtained from each participant prior to the experiment. The research was conducted with the ethical guidelines of the Faculty of Behavioural and Movement Sciences at the Vrije Universiteit Amsterdam and those laid down in the Declaration of Helsinki.

### Apparatus and stimuli

The experiment was run in a dimly lit cubicle and programmed using OpenSesame software (Mathot et al., 2012). Participants were located 57 cm from an LCD monitor (120 Hz refresh rate) and used a chinrest when performing the task. The stimulus display was presented on a black background (*<*.5 cd/m^2^). On each trial, three small red dots appeared on the screen (27 cd/m^2^). One of these, the fixation dot, was presented at the center of the screen. The other dots signified the possible target locations. The target was a green Gabor and centered on one of the two peripheral dot locations. Each target Gabor had a one-dimensional sinusoidal luminance profile, recruiting only the green component of the RGB display, with a spatial frequency of four cycles per degree of visual angle (d.v.a.) and a standard deviation of 0.25 d.v.a.. The green luminance modulation profile of each target Gabor ranged from a trough of *<*.5 cd/m^2^ (0, 0, 0) to a peak of either 48 cd/m^2^ (0, 255, 0; “bright” targets) or 14 cd/m^2^ (0, 145, 0; “dark” targets). Luminance contrasts were 95 and 27, respectively, using (Lmax – Lmin)/Lmin. The orientation of the target Gabor (orthogonal with respect to the orientation of its sinusoidal luminance profile) was 7 degrees counterclockwise or clockwise of vertical. Targets were presented either in isolation (i.e. target-alone condition) or in the context of two equidistant flanking distractors, one located above the target and the other below. On half of flanker-present trials, the flankers were both green (RGB: 0, 200, 0), with a maximum luminance of 31 cd/m^2^ (luminance contrast = 61) (Figures 1a, b); on the other half, they were both red (RGB: 200, 0, 0), with a maximum luminance of 14 cd/m^2^ (luminance contrast = 27) (Figures 1c, d). Of these green and red flanker conditions, both flanking Gabors were either vertically oriented (Figures 1a, c) or horizontally oriented (Figures 1b, d). Target– flanker distance was manipulated by varying the center-to-center separation of target and flanking Gabors (1, 2, 3, 4, 5, or 6 d.v.a.).

**Figure 1.**
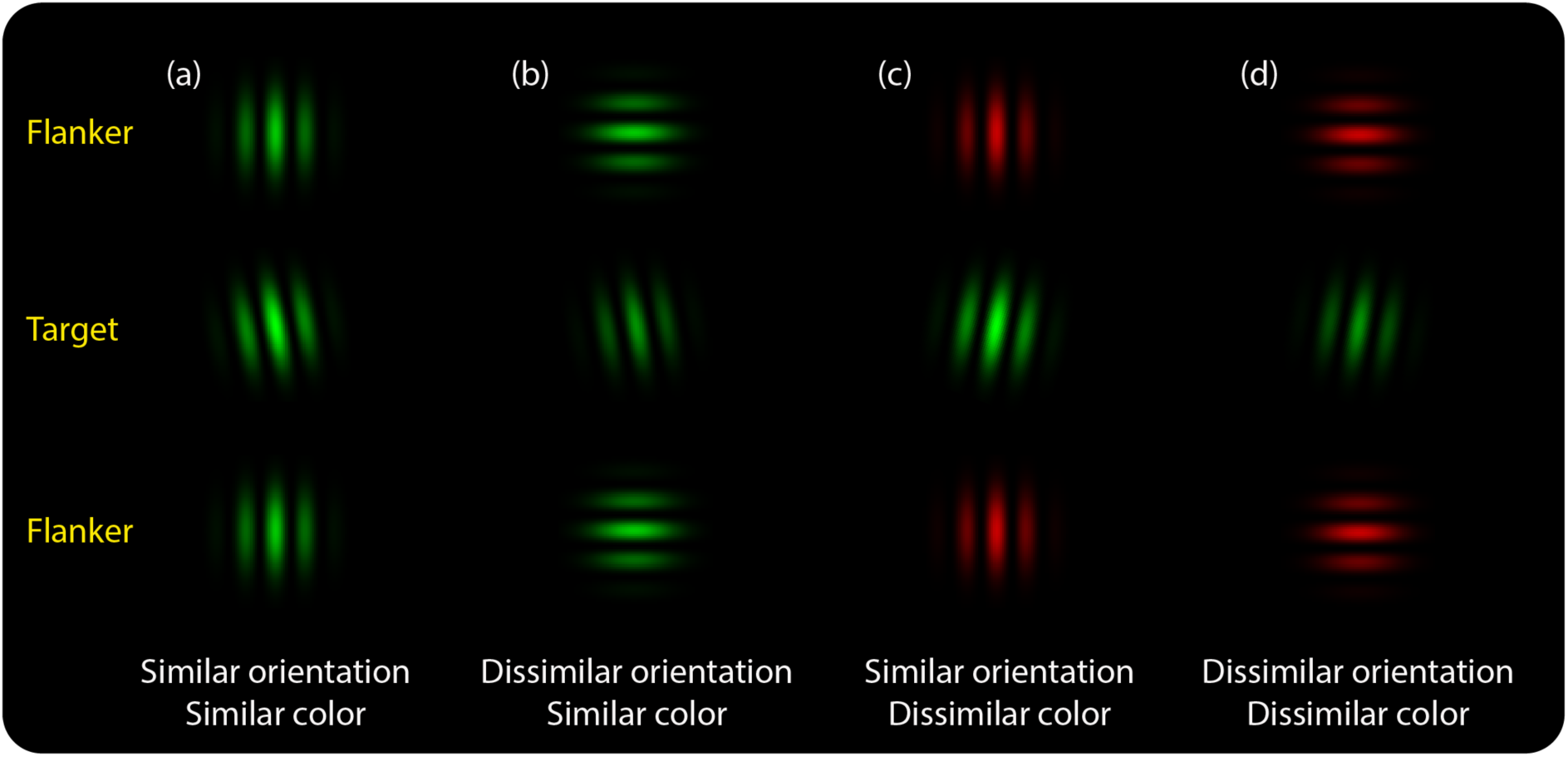
The four flanker feature combinations used in this study, in which target-relative color and/or orientation similarity is varied: (a) similar color, similar orientation; (b) similar color, dissimilar orientation; (c) dissimilar color, similar orientation; and (d) dissimilar color, dissimilar orientation. Note that in each trial, the target Gabor could be tilted either counter-clockwise (a & b) or clockwise (c & d) of vertical and be either low (b & d) or high luminance (a & c). Subjects performed either target luminance or orientation judgments in separate blocks of trials.

### Design and procedure

A trial started with the presentation of the three small red dots for 500 ms. Subsequently, a green target Gabor stimulus appeared on either the left or right side of fixation for 150 ms. This relatively short presentation interval was used to discourage participants from making eye-movements toward the target locations. Subjects were instructed to maintain fixation on the central red dot throughout the experiment. For the luminance discrimination task, participants were instructed to press the “d” or “l” key on a standard computer keyboard to signify whether the target appeared darker (“donker” in Dutch) or brighter (“lichter” in Dutch), respectively. For the orientation discrimination task, subjects were instructed to press the “z” or “m” (left or right) key if the peripherally presented target appeared tilted counterclockwise or clockwise of vertical, respectively. Participants were instructed that if they were unsure about the correct response on a given response that they should nonetheless choose a response based on their ‘best guess’. The subsequent trial was initiated when the participant made a response. Target location (left versus right), target orientation (–7° or +7° from vertical), target luminance (14 or 48 cd/m^2^), distractor orientation (same or different), distractor color (same or different), distractor distance (1, 2, 3, 4, 5, 6 d.v.a.), and a target alone condition were presented within blocks and selected randomly from trial to trial. The task order was counterbalanced across participants. For each of the two tasks, each distractor condition (green, red, vertical, or horizontal; each at six levels of separation) was presented 24 times, and the target-alone condition was presented 96 times. Each task consisted of 672 trials per participant and was divided into eight separate blocks of trials of 84 trials, separated by self-paced breaks.

## Results

### Brightness judgments

For all analyses, data from two participants were excluded as their performance was close to chance in the target-alone condition for one participant and below chance for the other (Cass & Van der Burg, 2023). Furthermore, practice trials as well as the first trial of each block were excluded from further analyses.

#### Serial dependence

First we examined whether the proportion of correct brightness judgements depended on the relative: target luminance (*t*-1 target luminance), target orientation (*t*-1 target orientation), and target location (*t*-1 target location) associated with the preceding trial *t*-1 compared to the current trial *t*. Figure 2 illustrates the proportion of correct brightness judgments as a function of the *t*-1 target location and *t*-1 target luminance when the target orientation on the preceding trial was either similar (panel A) or dissimilar to the current trial (panel B). The magnitude of serial dependence is depicted in the lower panels (i.e., *t*-1 similar brightness – *t*-1 dissimilar brightness).

**Figure 2.**
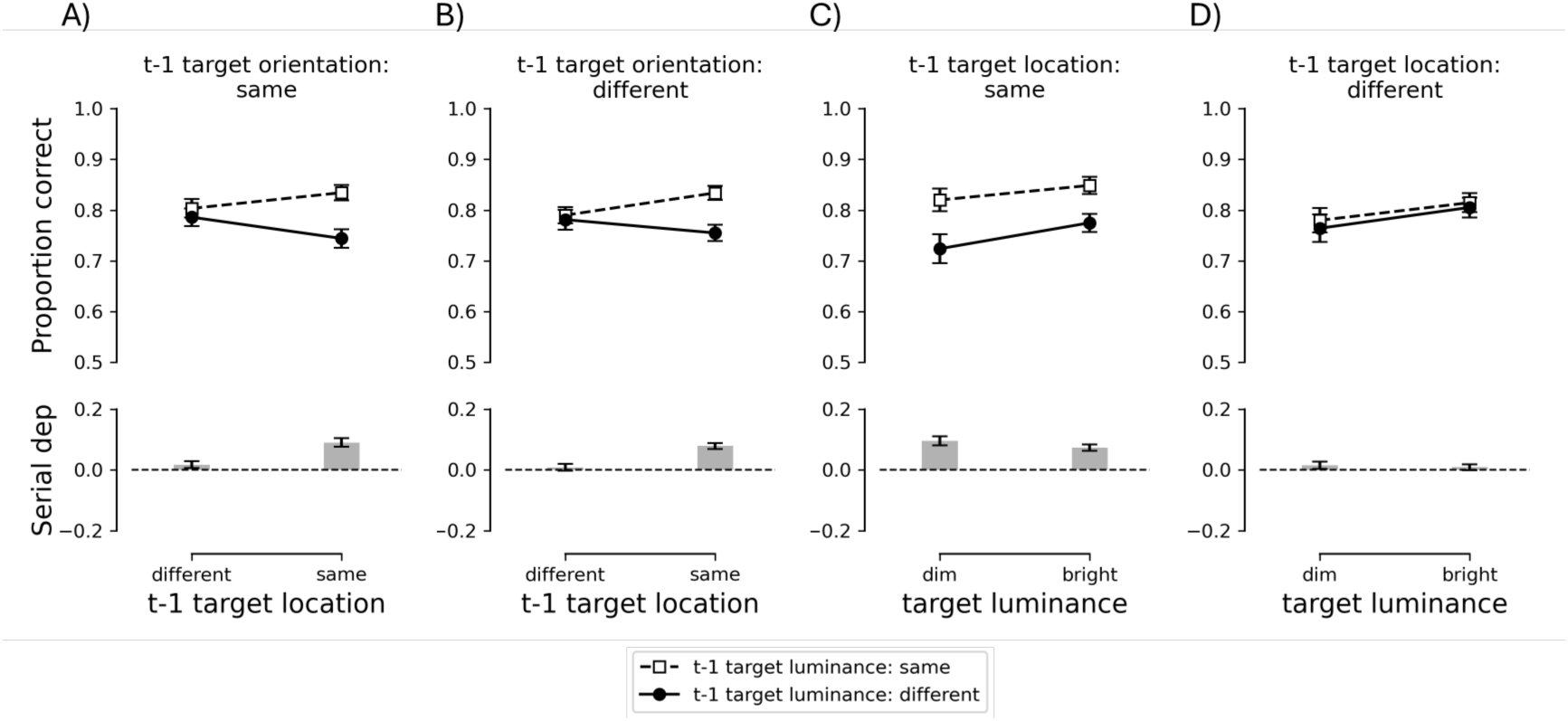
Serial dependence effects for the brightness judgment task. A and B) Proportion correct as a function *t*- 1 target location and *t*-1 target luminance when the target orientation on trial *t*-1 was similar (Panel A) or dissimilar to the target orientation on trial *t* (Panel B). C and D) Proportion correct as a function of the target luminance (on trial *t*) and the *t*-1 target luminance when the target location on *t*-1 was similar (Panel C) or dissimilar to the target location on trial *t* (Panel D). Lower panels reflect the serial dependence [i.e., *t*-1 target luminance: similar (dashed line) – *t*-1 target luminance: dissimilar (continuous line)]. Error bars represent between-subject mean standard errors.

We conducted a repeated-measures ANOVA on the mean proportion correct with *t*-1 target location (different or same), *t*-1 target luminance (same or different), and *t*-1 target orientation (same or different) as within subject factors. Alpha was set to .05. The ANOVA yielded a significant *t*-1 target luminance effect as well as a *t*-1 target luminance × *t*-1 target location interaction, *F*(1, 27) = 39.045, *p* < .001, ηp^2^ = .591, and *F*(1, 27) = 54.897, *p* < .001, ηp^2^ = .670, respectively. The two-way interaction was further examined using paired two-tailed *t*-tests for each *t*-1 target location. The *t*-test yielded a significant *t*-1 target luminance effect when the target on *t*-1 was presented at the same location as the current trial, *t*(27) = 8.216, *p* < .001, as the proportion correct was greater when the *t*-1 target luminance was similar (.83) than when it was dissimilar (.75; i.e., a positive serial dependence). In contrast, the *t*-test yielded no significant *t*-1 target luminance effect when the target was presented at a different location on the preceding trial, *t*(27) = 1.647, *p* = .111. The serial dependence brightness judgment effect did not depend on the *t*-1 target orientation, as all other effects were not significant, all *F*-values < 1.52, all *p*-values ≥ .228.

It is possible that the serial dependence brightness judgment effect described above could be driven by the specific target luminance values presented (dim or bright luminance). Therefore, we examined whether the observed serial dependencies themselves depended on the specific target luminance values presented on consecutive trials (see Figure 2c, and Figure 2d). We conducted a repeated measures ANOVA on the mean proportion correct with *t*-1 target location (different or same), *t*-1 target luminance (similar or dissimilar) and target luminance (dim or bright) as within-subject factors. The ANOVA yielded a significant *t*-1 target luminance effect as well as a two-way interaction between *t*-1 target luminance × *t*-1 target location, *F*(1, 27) = 39.376, *p* < .001, ηp^2^ = .593, and *F*(1, 27) = 62.379, *p* < .001, ηp^2^ = .698, respectively. The two-way interaction was further examined using two-tailed *t*-tests for each *t*- 1 target location. When the target was presented at the same location as the preceding trial (see Figure 2c), the proportion correct was significantly greater when the *t*-1 target luminance was similar to the target on trial *t* (.83) than when it was dissimilar (.75; i.e., a positive serial dependence), *t*(27) = 8.250, *p* < .001. In contrast, the *t*-test failed to reach significance when the target was presented at a different location compared to the preceding trial (see Figure 2d), *t*(27) = 1.664, *p* = .108. The serial dependence did not depend on the specific target luminance presented on consecutive trials, as all other effects failed to reach significance, all *F*-values < 1.801, all *p*-values ≥ .191.

When participants performed the brightness judgment task, we found a positive serial dependence for target luminance in a visual crowding paradigm. In other words, the proportion of correct brightness judgments was greater when the target on the preceding trial was similar to the current target than when it was dissimilar. Moreover, this effect was retinotopic, as the positive serial dependence was only observed when the target was presented at the same location as the target location on the preceding trial, and not when the target was presented at a different location. Furthermore, we show that the serial dependence neither depended on the target orientation on the current trial compared to the previous trial, nor on the specific target luminance on the current trial (i.e., serial dependencies occurred for both dim and bright targets).

#### Effects of visual crowding on the preceding trial on serial dependence

To assess whether visual crowding of brightness judgements in the preceding trial influenced serial dependence, we analyzed *t*-1 trials based on target-distractor distance. Visual crowding typically depends on both the similarity between target and distractors and their spatial proximity (Bouma, 1970; Cass & Van der Burg, 2023). Therefore, we excluded trials lacking a distractor in either trial *t*-1 or trial *t* and categorized the remaining data into two bins based on the target-distractor distance in trial *t*-1. Distances within Bouma’s window, where target-distractor separation was 1 or 2 d.v.a., were labeled as ’near,’ while those beyond Bouma’s window (5 or 6 d.v.a.) were labeled as ’far.’ Trials with intermediate distractor distances of 3 or 4 d.v.a. were excluded as they represent boundary conditions. Additionally, as our findings above indicate that the orientation of the *t*-1 target does not significantly influence serial dependence during brightness judgments, we disregarded target orientation in subsequent analyses of brightness judgments. Figure 3 illustrates the effect of visual crowding on the previous trial on both the magnitude and direction of serial dependence for the brightness judgment task.

**Figure 3.**
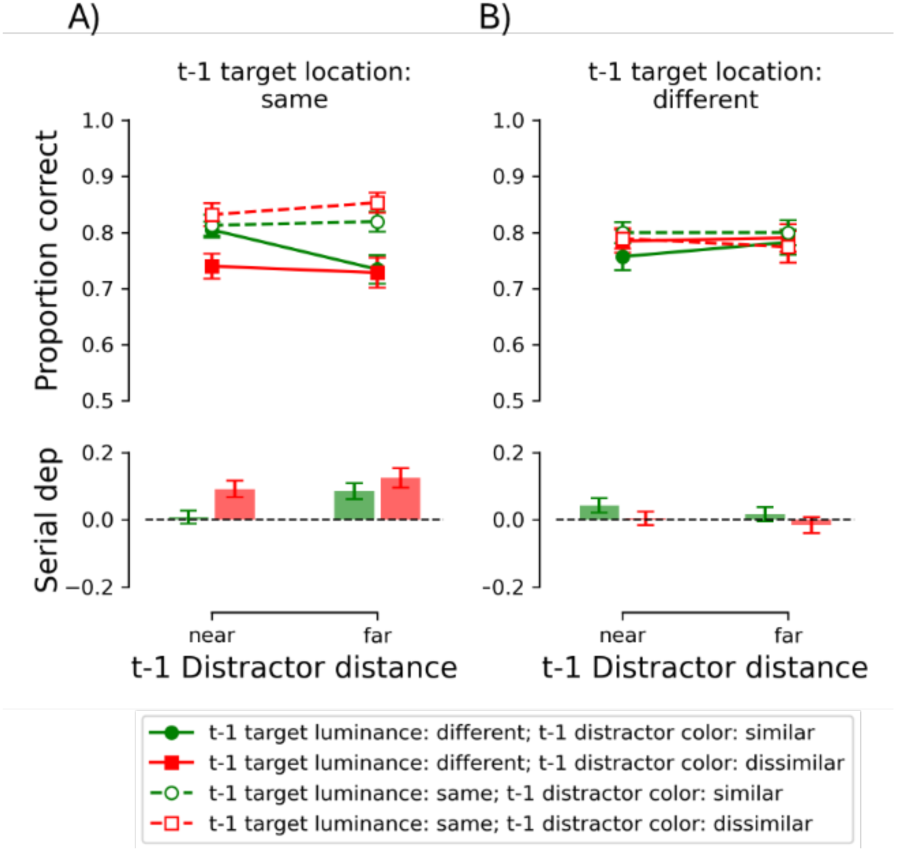
The effect of visual crowding on trial *t*-1 on serial dependence for the brightness judgment task. A) Mean proportion correct as a function of t-1 distractor distance (i.e., near: within Bouma’s window, or far: outside Bouma’s window), t-1 distractor color (compared to the target luminance) and t-1 target luminance for those trials wherein the target was presented at the same (Panel A) and different location as trial t-1 (Panel B). Lower panels reflect the serial dependence (i.e., *t*-1 target luminance: same – *t*-1 target luminance: different). Error bars represent between-subject mean standard errors.

We conducted an ANOVA on the proportion correct with *t*-1 distractor distance (near or far), *t*-1 distractor color (similar or dissimilar as the green target), *t*-1 target location (different or same) and *t*-1 target luminance (similar or dissimilar) as within-subject variables. The main effect of *t*-1 target luminance, the two-way interaction between *t*-1 target luminance and *t*-1 target location, and the three-way interaction among *t*-1 target luminance, *t*-1 target location *t*-1 distractor color were significant, all *F*-values ≥ 9.122, all *p*-values ≤ .005, all ηp^2^ ≥ .253. The three-way interaction was further examined by conducting two separate ANOVAs with *t*-1 target luminance and *t*-1 distractor color as within-subject variables for each *t*-1 target location. The ANOVA yielded no significant effects when the target on the preceding trial was presented at a different location compared to the current trial, all *F*-values < 2.222, all *p*-values ≥ .148 (see Figure 3b). In contrast, when the target was presented at the same location as the current trial (see Figure 3a), the *t*-1 target luminance × *t*-1 distractor color interaction was significant, *F*(1, 27) = 6.472, *p* = .017, ηp^2^ = .193. Two-tailed *t*-tests yielded a significant positive serial dependence when the distractors on the preceding trial were dissimilar to the target color, but also when the distractors were similar to the target color on the preceding trial, *t*(27) = 2.818, *p* = .009, and *t*(27) = 5.332, *p* < .001, respectively. Interestingly, the serial dependence magnitude was significantly greater when there was no crowding on trial *t*-1 (.108; collapse over the red bars in the Figure 3a) than when there was crowding on trial *t*-1 (.047; collapsed over the green bars in the Figure 3a), *t*(27) = 2.544, *p* = .017.

The three-way interaction among *t*-1 target luminance, *t*-1 target location and *t*-1 distractor distance was significant, *F*(1, 27) = 6.045, *p* = .021, ηp^2^ = .183. This interaction was further examined by conducting another ANOVA with *t*-1 target luminance and *t*-1 distractor distance for the same target location only. The ANOVA yielded a significant two-way interaction, *F*(1, 27) = 5.518, *p* = .0026, ηp^2^ = .170, indicating that the serial dependence effect was greater when the distractors were far away from the target on trial *t*-1 (.105) than when they were near the target (.050), *t*(27) = 2.349, *p* = .026. All other effects failed to reach significance, all *F*-values ≤ 3.944, all *p*-values ≥ .057, all ηp^2^ ≤ .127.

Visual crowding on the preceding trial, as determined by the distractors’ color (relative to the green target) as well as the distractor-target distance on trial *t*-1, had a significant influence on the magnitude of the serial dependence of brightness judgments, but only when the target was presented at the same location as on the preceding trial. Specifically, the serial dependence effect was greater when the stimuli were not crowded than when they were crowded on trial *t*-1.

#### Effect of serial dependence on visual crowding on the current trial

We examined whether serial dependence has an effect on visual crowding on the current trial. Like the previous analyses, we excluded trials when there was no distractor present on trial *t* or *t*-1, and we segregated the data into two bins, wherein the target-distractor distance was within Bouma’s window (i.e., the target-distractor distance was 1, or 2 d.v.a.; labelled as ‘near’) or outside Bouma’s window (i.e., the target-distractor distance was 5 or 6 d.v.a.; labelled as ‘far’). Trials on which the distractor distance on trial *t* was either 3 or 4 d.v.a. were excluded as they belong to the boundary conditions. Figure 4 illustrates the effect of serial dependence on visual crowding for the brightness judgment task.

**Figure 4.**
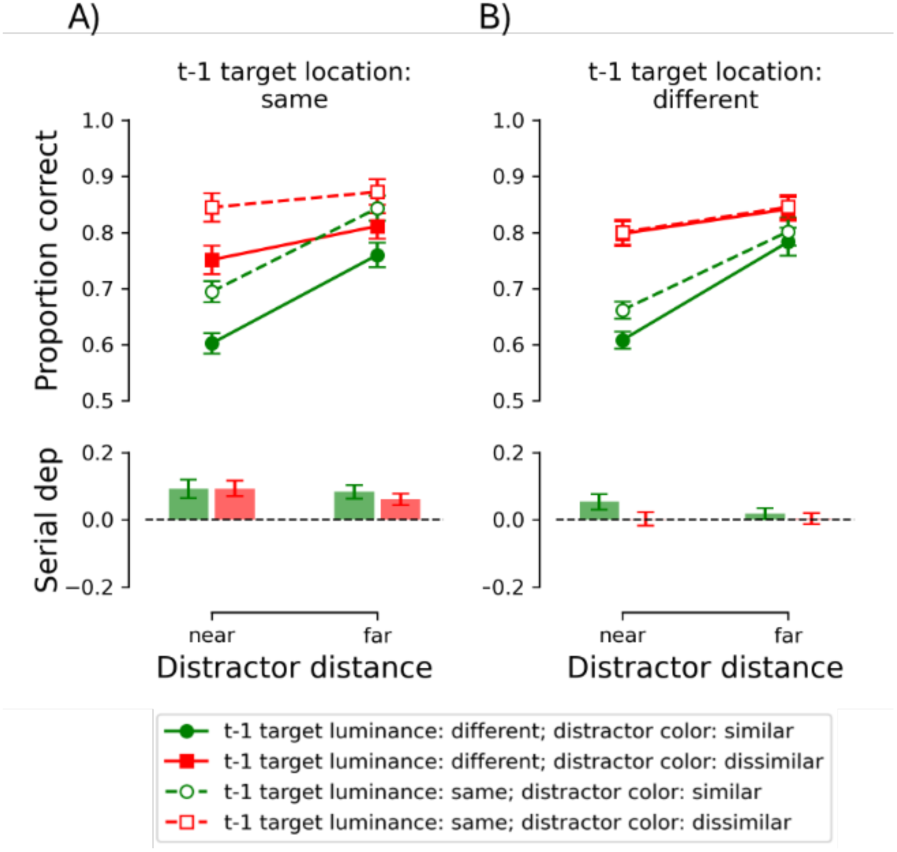
The effect of serial dependence on visual crowding on the current trial for the brightness judgment task. A) Mean proportion correct as a function of distractor distance (i.e., near: within Bouma’s window, or far: outside Bouma’s window), distractor color (compared to the target color), and *t*-1 target lumiance for trials in which the target was presented at the same location as trial *t*-1 (Panel A) or at a different location (Panel B). Lower panels reflect the serial dependence (i.e., *t*-1 target luminance: similar – *t*-1 target luminance: dissimilar). Error bars represent between-subject mean standard errors.

We conducted an ANOVA on the proportion correct with distractor distance (near or far), distractor color (similar or dissimilar as the green target), *t*-1 target location (different or same) and *t*-1 target luminance (similar or dissimilar) as within subject variables. The main effect of *t*-1 target luminance, distractor color and distractor distance were significant, all *F*- values ≥ 30.969, all *p*-values < .001. Consistent with our previous analysis, the *t*-1 target luminance × *t*-1 location was significant, *F*(1, 27) = 34.955, *p* < .001, ηp^2^ = .564, as the positive serial dependence was retinotopic (see previous analyses). The two-way interaction between distractor color and distractor-target distance was significant, *F*(1, 27) = 30.749, *p* < .001, ηp^2^ = .532, as the proportion correct depended on both the distractor color (i.e., target-distractor similarity) and the distance between the target and the distractor (i.e., visual crowding). All other effects failed to reach significance, *F*-values ≤ 2.981, all *p*-values ≥ .096, indicating that visual crowding did not depend on serial dependence.

Although we observed a positive serial dependence when participants made brightness judgments, this did not affect visual crowding (i.e. the effects of target-flanker distance and target-flanker color similarity). More specifically, the serial dependence effect had an additive effect on brightness judgment performance.

### Orientation judgments

#### Serial dependence

Figure 5 illustrates the proportion correct as a function of the target location on *t*-1, the target orientation on *t*-1 when the target luminance on trial *t*-1 was similar to the current trial (panel A) and dissimilar to the current trial (panel B). The serial dependence magnitude is depicted in the lower panels (i.e., *t*-1 similar orientation – *t*-1 dissimilar orientation).

**Figure 5.**
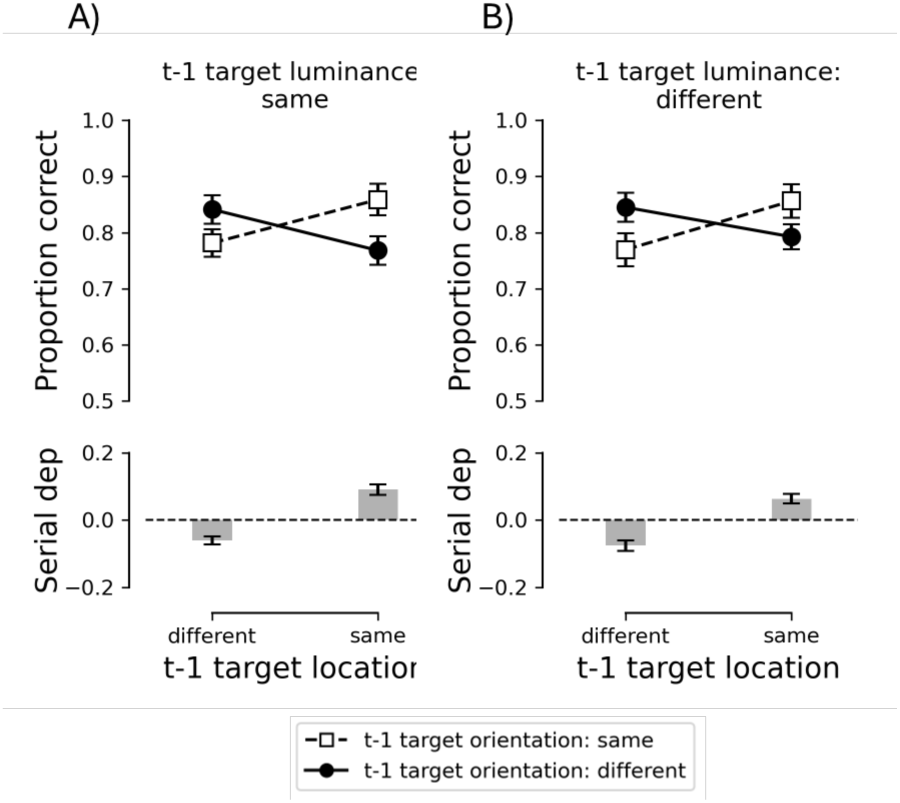
Orientation judgment results. A) Mean proportion correct as a function of *t*-1 target location, *t*-1 target orientation when the *t*-1 target luminance was similar (Panel A) or dissimilar (Panel B) to the current target luminance. Lower panels reflect the serial dependence (i.e., *t*-1 target orientation: similar – *t*-1 target orientation: dissimilar) as a function of the target location on trial *t*-1. Error bars represent between-subject mean standard errors.

We conducted an ANOVA on the mean proportion correct with *t*-1 target location, *t*-1 target orientation, and *t*-1 target luminance as within subject variables. The ANOVA yielded a significant *t*-1 target orientation × *t*-1 target location interaction, *F*(1, 27) = 75.475, *p* < .001, ηp^2^ = .737. The interaction was further examined using two-tailed *t*-tests for each *t*-1 target location. When the previous target location was the same as the current trial, the proportion correct was significantly greater when the target orientation was similar to the preceding trial (.86) than when it was dissimilar to the preceding trial (.78; i.e., a positive serial dependence), *t*(27) = 7.127, *p* < .001. When the previous target location was different to the current trial, the proportion correct was significantly lower when the target orientation was similar to the preceding trial (.78) than when it was dissimilar to the preceding trial (.84; i.e., a negative serial dependence), *t*(27) = 5.236, *p* < .001. The ANOVA yielded no other significant effects, *F*-values ≤ 2.985, all *p*-values ≥ .095, all ηp^2^ ≤ .100.

Taken together, we found a significant serial dependence effect when participants made orientation judgments. This serial dependence was retinotopic, and did not depend on the specific target luminance present on the preceding trial. More specifically, a positive serial dependence was observed when the target was presented at the same location as the preceding trial, and a negative serial dependence was observed when the target was presented at the opposite (i.e., different) location on the preceding trial.

#### Effect of visual crowding on the preceding trial on serial dependence

Next, we examined whether the effect of visual crowding on the previous trial had a significant effect on serial dependence. Like in our previous analyses for brightness judgments, we excluded trials when there was no distractor present on trial *t*-1 or trial *t*, and split the data into two bins: inside (i.e., the target-distractor distance was 1 or 2 d.v.a., labelled as ‘near’) or outside Bouma’s window (i.e., the target-distractor distance was 5 or 6 d.v.a., labelled as ‘far’). Furthermore, given that the *t*-1 target luminance has no significant effect on serial dependence (when performing an orientation judgment), we decided to ignore the target luminance (dim or bright) for the following analyses. Figure 6 illustrates the effect of visual crowding on the previous trial on serial dependence for the orientation judgment task.

**Figure 6.**
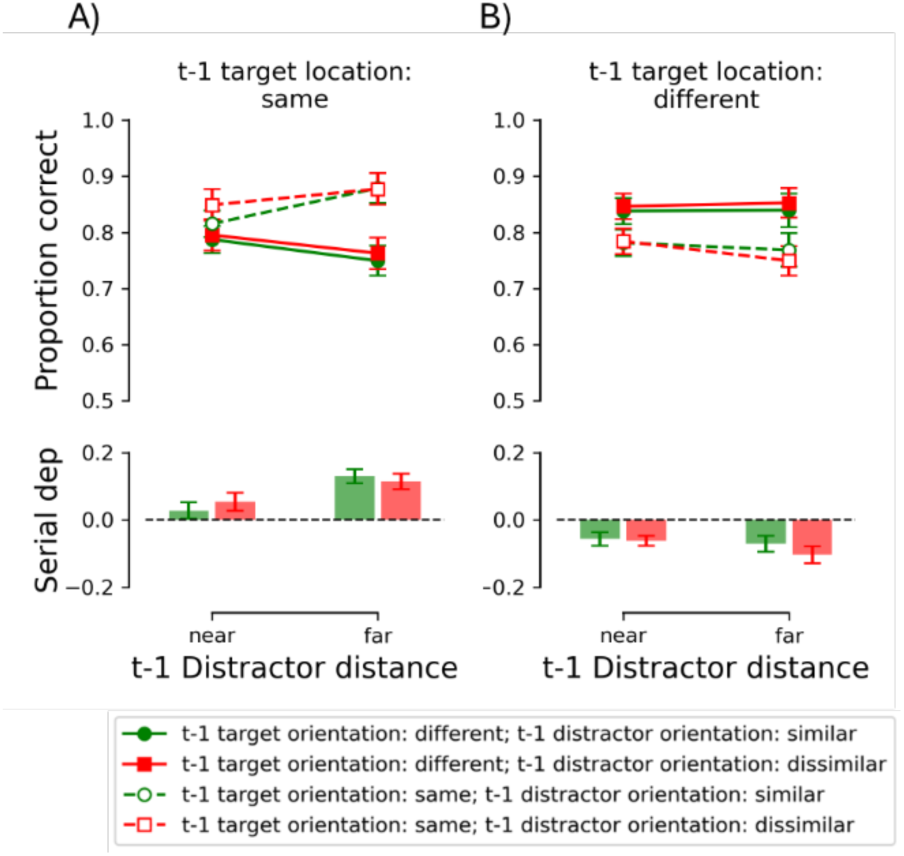
The effect of visual crowding on trial *t*-1 on serial dependence for the orientation judgment task. Mean proportion correct as a function of *t*-1 distractor distance (i.e., near: within Bouma’s window, or far: outside Bouma’s window), *t*-1 distractor orientation (compared to the nearly vertical target) and *t*-1 target orientation for those trials in which the target was presented at the same (Panel A) or different location (Panel B) compared to the current trial. Lower panels reflect the serial dependence (i.e., *t*-1 target orientation: similar – *t*-1 target orientation: dissimilar). Error bars represent between-subject mean standard errors.

We conducted an ANOVA on the mean proportion correct with *t*-1 distractor distance, *t*-1 distractor orientation (compared to the *t*-1 near vertical target), *t*-1 target orientation and *t*- 1 location as within subject variables. The ANOVA yielded a significant *t*-1 location × *t*-1 target orientation interaction (*F*(1, 27) = 81.787, *p* < .001, ηp^2^ = .752), a *t*-1 target orientation × *t*-1 distractor distance interaction (*F*(1, 27) = 4.764, *p* = .038, ηp^2^ = .150), and a significant *t*- 1 location × *t*-1 target orientation × *t*-1 distractor distance interaction (*F*(1, 27) = 10.292, *p* = .003, ηp^2^ = .276). The three-way interaction was further examined by conducting an ANOVA with *t*-1 target orientation and *t*-1 distractor distance as within subject variables for each *t*-1 location. When the target was presented at the same location as on the preceding trial, the ANOVA yielded a significant interaction, *F*(1, 27) = 12.826, *p* = .001, ηp^2^ = .322, indicating a smaller positive serial dependence when the distractors on trial *t*-1 were presented within close proximity of the target (.041) than when they were more distant from the target (.122). Note that the serial dependence (i.e., accuracy when *t*-1 target orientation was similar to the current target orientation – accuracy when *t*-1 target orientation was dissimilar) was significant for both distractor distances on trial *t*-1, *t*(27) ≥ 2.479, *p* ≤ .02. When the target was presented at the opposite location on the preceding trial, the ANOVA yielded a significant *t*-1 target orientation effect, *F*(1, 27) = 46.182, *p* < .001, ηp^2^ = .631, reflecting the negative serial dependence. The main effect of distractor distance on *t*-1 and the two-way interaction failed to reach significance, *F*-values ≤ 2.126, all *p*-values ≥ .158, all ηp^2^ ≤ .073, indicating that the serial dependence did not depend on crowding on the preceding trial when the target was presented at different visual locations on subsequent trials. The main ANOVA yielded no other significant effects, *F*-values ≤ 1.777, all *p*-values ≥ .194, all ηp^2^ ≤ .062.

Crowding on the preceding trial only had a significant effect on serial dependence when the target was presented at the same location as on the current trial. More specifically, the positive serial dependence was significantly reduced when the distractors in trial *t*-1 were presented in close proximity to the target location, compared to when the distractors were positioned further from the target in trial *t*-1. The distractor orientation on the preceding trial had no significant effect on the serial dependence. A negative orientation serial dependency was observed when target was presented in the opposite visual field across subsequent trials. This negative serial dependency was unaffected by target-distractor proximity (crowding) on trial *t*-1.

#### Effect of serial dependence on visual crowding on the current trial

We investigated whether serial dependence affects visual crowding in the current trial. To this end, we excluded trials in which no distractor was present in either trial *t* or trial *t*-1, and divided the data into two bins (i.e., the target-distractor distance was 1 or 2 d.v.a., labelled as ‘near’) or outside Bouma’s window (i.e., the target-distractor distance was 5 or 6 d.v.a., labelled as ‘far’). Figure 7 illustrates the effect of serial dependence on visual crowding for the orientation judgment task.

**Figure 7.**
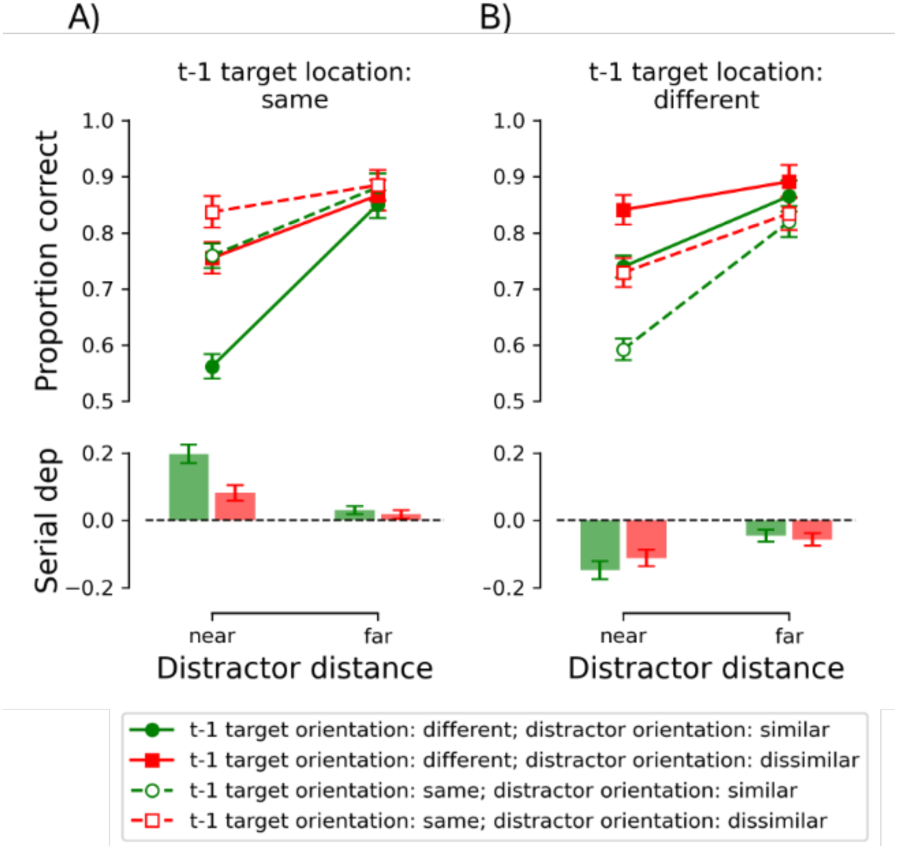
The effect of serial dependence on visual crowding for the orientation judgment task. A) Mean proportion correct as a function of distractor distance (i.e., near: within Bouma’s window, or far: outside Bouma’s window), distractor orientation (compared to the target orientation) and *t*-1 target orientation when the target was presented at the same location (Panel A) or at a different location (Panel B) compared to trial *t*-1. Lower panels reflect the serial dependence (i.e., *t*-1 target orientation: similar – *t*-1 target orientation: dissimilar). The error bars represent the sem.

We conducted an ANOVA on the mean proportion correct with *t*-1 target location, *t*-1 target orientation, distractor distance and distractor orientation as within-subject variables. The ANOVA yielded a significant four-way interaction, *F*(1, 27) = 8.580, *p* = .007, ηp^2^ = .241. We further examined this interaction using two separate ANOVAs with *t*-1 target orientation, distractor distance, and distractor orientation as within subject variables for each *t*-1 target location.

When the target location was the same as on trial *t*-1 (see Figure 7a), the ANOVA yielded a significant three-way interaction, *F*(1, 27) = 13.834, *p* < .001, ηp^2^ = .339, indicating that serial dependence (i.e., the *t*-1 target orientation) had an effect on visual crowding (i.e., the distractor distance × distractor orientation interaction). The magnitude of visual crowding was assessed for both *t*-1 target orientations by calculating the difference between conditions where the distractors were similarly oriented to the target and where the distractors had a dissimilar orientation on the current trial. When the distractors were presented close to the target on the current trial, the visual crowding magnitude (performance correct for near vs far distractor conditions) was significantly larger when the target orientation on the preceding trial was dissimilar (.19; see the continuous lines in Figure 7a) than when the target on the preceding trial was similar to the target on the current trial (.08; see the dashed lines in Figure 7a), *t*(27) = 4.558, *p* < .001. Conversely, when the distractors were positioned far from the target in the current trial, the magnitude of visual crowding remained unaffected by the orientation of the target in trial *t*-1., *t*(27) = .790, *p* = .436.

When the target location was different to the *t*-1 target location (see Figure 7b), the ANOVA yielded a significant distractor distance × distractor orientation interaction, *F*(1, 27) = 17.668, *p* < .001, ηp^2^ = .396, indicating a significant visual crowding effect. This crowding effect did not depend on the *t*-1 target orientation, as the three-way interaction failed to reach significance, *F*(1, 27) = 1.353, *p* = .255, ηp^2^ = .048. However, the *t*-1 target orientation interacted with the distractor distance, *F*(1, 27) = 15.266, *p* < .001, ηp^2^ = .361, with serial dependence being stronger when distractors were positioned close to the target compared to when they were positioned farther away (which might be explained by a ceiling effect in the latter condition). All main effects were significant, *F* values ≥ 27.278, *p* values < .001, ηp^2^ ≥ .503. The *t*-1 target orientation × distractor orientation analysis failed to reach significance, *F*(1, 27) = .283, *p* = .599, ηp^2^ = .010.

In summary, during the orientation task, visual crowding was influenced by serial dependence when the target appeared at the same location as in the previous trial, but not when presented at the opposite location. In the former case, visual crowding increased when the target orientation on a given trial differed from that in trial *t*-1, compared to when they were similar.

### Correlations between visual crowding and serial dependence effects

Finally, we examined whether visual crowding and serial dependencies for both brightness as well as orientation judgments correlated. Figure 8 illustrates the correlation between visual crowding and serial dependencies for both judgments. Note that for the serial dependencies, the data was divided into t-1 target location same versus different as the serial dependencies were retinotopic specific.

**Figure 8.**
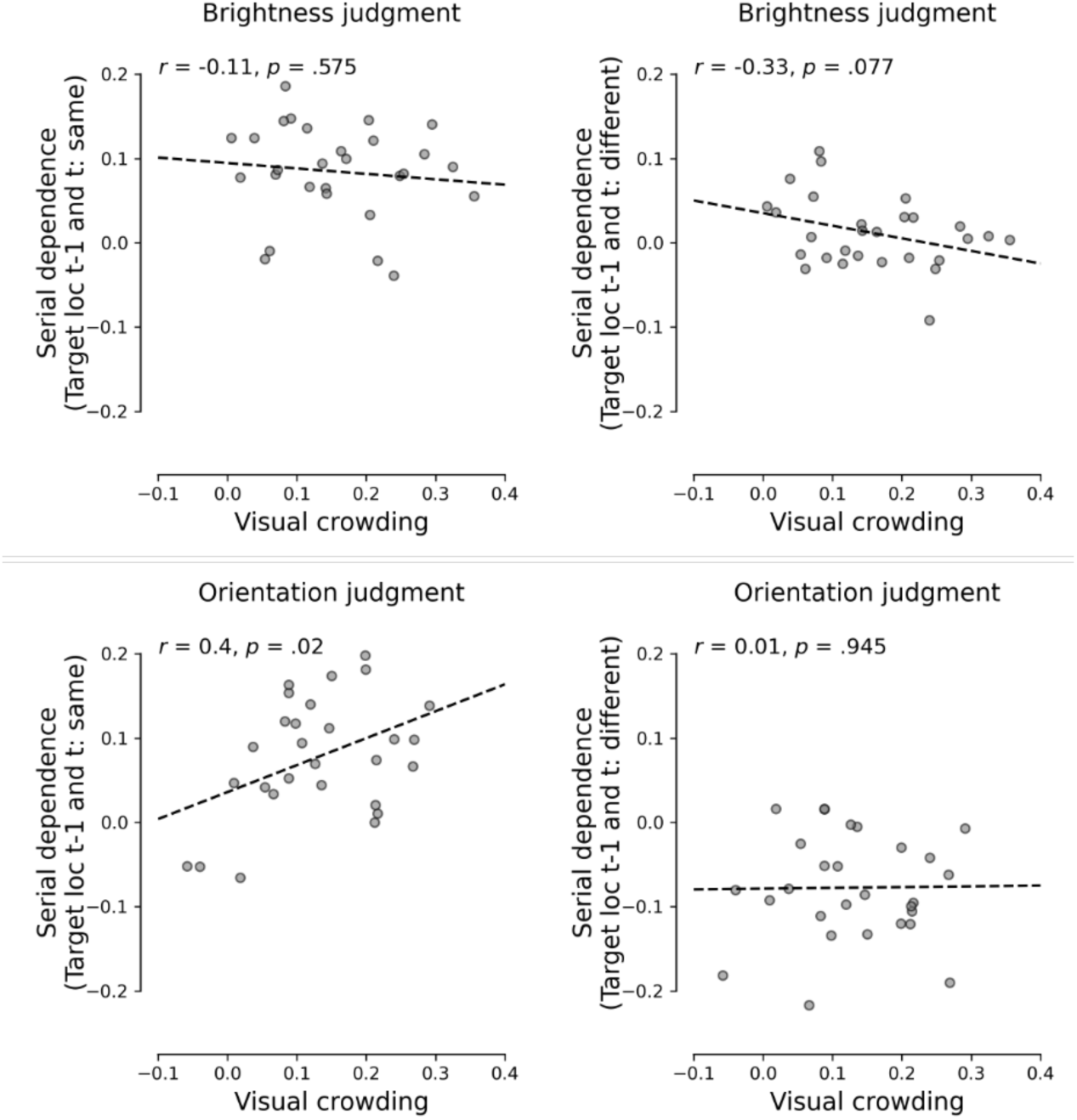
Correlations between serial dependence and visual crowding for brightness (upper panels) and orientation judgments (lower panels). For the left panels, for the serial dependence only, the target was presented at the same location compared to the target location on the preceding trial. For the right panels, the target was presented at the different location compared to the previous trial. Serial dependence was defined by subtracting the proportion correct for similar targets on two consecutive trials - the proportion correct for dissimilar targets on two consecutive trials. Visual crowding was defined by subtracting the proportion correct for different nearby distractors (i.e., red distractors for brightness judgments, and horizontal distractors for orientation judgments) - the proportion correct for similar nearby distractors (i.e., green distractors for brightness judgments, and vertical distractors for orientation judgments). Pearson correlations are shown in each panel.

For the brightness (luminance) judgments, the analyses yielded no significant correlations between visual crowding and serial dependence (when the target location was the same as on the preceding trial: Pearson *r* = -.11, *p* = .575; when the location was different: Pearson *r* = -.33, *p* = .077). In contrast, the analyses yielded a significant correlation between visual crowding and serial dependencies when the target location was the same as on the preceding trial (Pearson *r* = .40, *p* = .02), but not when it was different (Pearson *r* = .01, *p* = .945).

## Discussion

This study examined the functional relationship between serial dependence and visual crowding in both orientation and brightness judgments using identical stimuli (in randomised sequences) for both tasks. We first determined whether serial dependence and crowding share functional similarities, particularly in terms of feature-specific dependencies related to perceptual task demands. Secondly, we investigated the causal structure of serial dependence and crowding by exploring their reciprocal interactions. Finally, we assessed whether the magnitudes of these effects were correlated for each task type. This exploration helped clarify the functional relationship of serial dependence and visual crowding (if any) and explore how spatial and temporal visual integration processes might coordinate within a unified predictive framework.

Our findings provide evidence that serial dependence and visual crowding are functionally linked. Specifically, the stimulus properties driving these effects are task-specific: brightness judgments are linked to relative target luminance (not orientation), while orientation judgments are linked to relative target orientation (not luminance). This double dissociation suggests that the mechanisms underlying serial dependence for brightness and orientation judgments are likely mediated by independent neural processes. A similar dissociation was observed in our prior crowding study, where brightness judgments depended on target/flanker color similarity, while orientation judgments were driven by target/flanker orientation (Cass & Van der Burg, 2023). Similarly, brightness judgments depended upon target/flanker colour similarity, not flanker orientation, and vice versa for orientation judgments. This striking similarity in the task and stimulus features that govern serial dependence and visual crowding effects supports the hypothesis that these phenomena may be functionally linked.

While we show that both visual crowding (Cass & Van der Burg, 2023) and serial dependence processes each exhibit feature-specific independence that depends on the task at hand, they interact in a structured manner, predominantly influenced by spatial context in the case of visual crowding. This aligns with predictive coding theories, such as those proposed by Cicchini et al. (2022), which suggest that serial dependence and crowding represent complementary predictive coding strategies for leveraging temporal and spatial image redundancies. Our findings support this hypothesis, but also reveal specific boundary conditions. For example, we found that crowding occurring on the immediately preceding trial reduces the magnitude of (positive) serial dependence, but only when the target appears in the same location.

In predictive coding terms, crowding functions as a spatial ’gatekeeper,’ determining whether temporal smoothing via serial dependence should be applied. In crowded conditions, where spatial redundancy is high and individual features are difficult to resolve, the brain prioritizes immediate spatial context over prior visual experiences. This modulation of serial dependence by crowding suggests a hierarchical approach to redundancy reduction: spatial integration precedes temporal predictions, with serial dependence engaged only when spatial clarity allows. Consequently, the weakening of serial dependence under high crowding conditions may reflect an adaptive mechanism to avoid over-assimilating ambiguous visual information. An alternative, simpler explanation—but also compatible with predictive coding—is that crowding disrupts the encoding of previous target stimulus features, preventing the formation of a robust foundation for serial dependence. The distinction lies in whether the weight of prior information is dynamically adjusted (as predictive coding suggests) or whether inadequate encoding of previous stimuli (due to crowding) undermines the effect altogether.

Our findings also reveal a task-dependent asymmetry. For brightness judgments, while crowding constrains serial dependence, serial dependence does not reciprocally influence crowding. In contrast, for orientation judgments, serial dependence interacts directly with crowding, amplifying or mitigating its effects based on the spatial and temporal similarity of stimuli. This interaction underscores the dynamic nature of orientation judgments, where serial dependence and crowding combine to shape perception depending on spatio-temporal contextual conditions. Moreover, the observed relationship for orientation judgments—where stronger serial dependence corresponds to greater crowding on the current trial—again highlights the dominance of immediate spatial conditions over prior experiences in cluttered environments. Together, these findings suggest that while both crowding and serial dependence may contribute to predictive coding (redundancy reduction), they do so with distinct task-specific functional roles and varying degrees of influence. The observation that stronger positive serial dependence is associated with greater crowding on the current trial underscores the dominant role of current spatial stimulus conditions in visual perception, particularly in cluttered environments. The observation that crowding affects serial dependence but not vice versa in brightness judgments further supports the notion that spatial contexts predominantly govern temporal contexts (serial dependence), reinforcing the visual system’s prioritization of immediate spatial conditions over prior temporal input in cluttered visual environments.

Our correlational analysis revealed another facet to the relationship between crowding and serial dependence, which was again highly task-specific. The significant positive correlation between the magnitude of these effects for orientation judgments in which the target appeared in the same location as the previous trial is indicative of an even more direct connection between them. That being said, the moderate strength of the correlation (*r* = 0.4) implies that while they may overlap, it is unlikely that they depend upon entirely identical neural mechanisms. Future research using neuroimaging techniques could clarify the degree to which these processes may or may not recruit similar neural resources to accomplish their complementary and overlapping functions. Moreover, the fact that even for the orientation judgments, the correlation depended upon correspondence of target locations across trials further illustrates the specificity of these interactions.

A key difference between the serial dependencies observed for brightness and orientation judgments is the presence of robust negative serial dependencies for the latter judgments when the target (and distractors) are located at different visual locations (opposite visual fields) across consecutive trials. Specifically, in these cases, participants judged target orientations more accurately when the target’s orientation differed across trials, but only when the target appeared in the opposite visual field. Negative serial dependencies have been observed in previous studies of orientation judgments, with multiple studies showing similar effects when the target appears in different locations across trials (Manassi et al., 2023) (but see Fritsche et al. (2017) who found that repulsive serial dependencies for orientation judgments are retinotopically specific). This repulsive serial dependence effect observed for orientation judgments at different retinal location suggests a strategy that enhances sensitivity to changes over time, particularly when faced with spatially disparate oriented stimuli. This may be mediated by long-range orientation facilitation or disinhibition (Angelucci et al., 2002; Walker et al., 2002), allowing the visual system to adaptively prioritize subtle variations in orientation across the visual field, possibly signalling the presence of a new object. The absence of a similar effect in brightness judgments indicates that the system treats temporal continuity in these domains differently—orientation judgments benefit from a dynamic adjustment to spatial shifts, while brightness judgments rely on a more stable integration of information across time. These contrasting directions of serial dependence effects reveal the manner in which the brain is able to exploit prior information not only to retain stable representations of its environment, but also to detect change.

What are the functional implications of these differences in how serial dependence and visual crowding interact across brightness and orientation task domains? First, consider the non-reciprocal interaction between visual crowding and serial dependence in brightness judgments. In natural vision, retinal luminance variations due to reflected light are principally caused by both illumination changes (e.g., shadows, dappled light) and the reflectance properties of object surfaces. Cass and Van der Burg (2023) showed that brightness judgments are modulated by visual crowding, depending on the hue similarity between the target and flankers. Significant brightness crowding occurred when the target and flankers shared the same hue (e.g., green), but crowding effects diminished or disappeared when their hues differed (e.g., red vs. green). They propose that this hue-specific crowding may serve an adaptive purpose: reducing perceptual representations of luminance variations caused by changing illumination (e.g., shadows in foliage) while preserving reflectance-based signals (e.g., hue contrast like red-green) (Cass & Van der Burg, 2023). Such a mechanism would allow the visual system to focus its processing resources on object segmentation (rather than illumination variation) in cluttered natural environments. Given this, the finding that serial dependence does not reciprocally influence crowding in brightness tasks may reflect a functional advantage. Because illumination variations (e.g. shadows) can change rapidly, historical information may be unreliable for brightness judgments, leading the visual system to prioritize immediate spatial signals over temporal smoothing to maintain accurate representations of surface reflectance.

Now, consider the reciprocal interaction between serial dependence and crowding in orientation judgments, where crowding on a prior trial reduces serial dependence, and stronger serial dependence on the current trial enhances crowding. Orientation information is critical for detecting edges, contours, and surface boundaries—key components of object segmentation, scene organization, and navigation. Crowding disrupts the perception of edges and contours by integrating/grouping of spatially proximate signals (Dakin et al., 2010; Greenwood et al., 2010; Herzog et al., 2015; Parkes et al., 2001; Van der Burg, Cass, et al., 2024), while serial dependence smooths orientation information over short time windows (Fischer & Whitney, 2014). However, when crowding on a prior trial reduces serial dependence, it may represent a recalibration mechanism to prevent excessive reliance on prior stimuli under conditions of strong spatial interference. Conversely, when serial dependence is strong, current crowding effects intensify, increasing spatial integration and amplifying perceptual contamination by nearby features. This reciprocal modulation may reflect a dynamic ecological strategy: unlike brightness judgments, where the system minimizes contamination from prior stimuli to prioritize reflectance-based clarity, orientation judgments rely more on the balance between past and present signals. The observed interactions demonstrate how serial dependence and crowding may work together to adaptively prioritize spatial and temporal conditions for optimal edge detection and object segmentation in cluttered visual environments.

Although further research is required to pinpoint specific neural loci for the temporal and spatial contextual effects observed in this study, the known emergence of orientation selectivity in the primary visual cortex (V1: (Boynton, 2005; Hubel & Wiesel, 1962, 1968)) strongly supports a cortical locus for both serial dependence and visual crowding as they pertain to orientation judgments. In contrast, the mechanisms underlying the positive serial dependence observed in brightness judgments are less clear, as luminance is encoded at the earliest stages of retinal processing. Notably, our observation that brightness serial dependencies are modulated by crowding effects on immediately preceding trials implies that the neural mechanisms responsible for brightness-based serial dependence likely involve input from neurons beyond the classical receptive fields associated with V1. Therefore, we propose that orientation and brightness (positive) serial dependencies are mediated by cortical visual mechanisms, possibly involving inter-blob and blob regions (Livingstone & Hubel, 1984) in V1 and/or thin and thick stripe regions in V2 (Roe & Ts’o, 1995; Ts’o & Gilbert, 1988).

A final point we would like to make concerns the task-stimulus contingencies observed here. Similar double dissociations between task and stimulus dependencies to those observed in this study and Cass and Van der Burg (2023) have been observed in visual crowding paradigms for other tasks. For example, Greenwood and Parsons (2020) and Yashar et al. (2019) each reported a double dissociation between task and visual stimulus properties (color-motion and orientation-color, respectively). Conversely, some stimulus-task combinations, such as orientation and spatial frequency (Yashar et al., 2019) and position and orientation (Greenwood et al., 2012), show non-independence during visual crowding. Understanding how these task and stimulus dimensions interact over time, particularly in relation to serial dependence, is crucial for determining whether the spatio-temporal redundancy reduction principles inferred from the current study apply more broadly. We encourage future crowding studies to address the existence and influence of temporal sequential effects of the kind revealed here, focusing on the functional and potentially causal relationship between crowding and serial dependence in other tasks and visual stimulus domains. These interactions could shed fresh explanatory light on previous findings in the crowding literature and refine our understanding of the underlying mechanisms.

## Conclusion

This study is the first to investigate the functional relationship between serial dependence and visual crowding, providing new insights into their interactions and underlying mechanisms. Both phenomena exhibit strikingly similar functional double dissociations: brightness judgments depend on relative luminance but not orientation, while orientation judgments are driven by orientation similarity but unaffected by luminance. This parallel dissociation suggests distinct but functionally linked neural mechanisms that support spatial and temporal integration in a task-dependent manner.

Our findings highlight task-specific interactions between serial dependence and visual crowding. For orientation judgments, crowding diminishes serial dependence under conditions of high spatial redundancy, while serial dependence amplifies crowding effects, indicating a reciprocal relationship. For brightness judgments, crowding constrains serial dependence without reciprocal effects, reflecting a prioritization of spatial integration/grouping over temporal smoothing. Positive serial dependencies—and their interactions with crowding—are retinotopically specific, occurring only when the target appears in the same location across consecutive trials. In contrast, negative serial dependencies, observed only for orientation judgments, emerged exclusively when the target’s location changed between trials. These findings suggest that positive serial dependencies may enhance temporal stability within consistent spatial contexts, while negative dependencies could promote sensitivity to spatially disparate changes, particularly for orientation processing, which is crucial for edge detection and object segmentation.

Additionally, the positive correlation between crowding and serial dependence for orientation judgments—but not brightness—indicates closer coupling for orientation, possibly reflecting overlapping neural pathways. The presence of robust negative serial dependencies when target locations changed further suggests that orientation judgments may leverage long-range mechanisms to adaptively respond to spatial variation, in contrast to brightness judgments’ reliance on stable spatial integration.

Taken together, these findings are consistent with the idea that the brain dynamically balances spatial and temporal inputs to optimize perceptual outcomes depending on task demands and stimulus characteristics. The shared double dissociations, spatially specific positive serial dependencies, and location-dependent negative serial dependencies underscore the complexity of these interactions and their potential contribution to efficient visual processing. Future research should explore the neural substrates of these processes as well as their broader implications for other visual and multisensory task and feature domains.

